# Water hardness variously influences tissue physiology of freshwater fish - *Cyprinus carpio* var *koi*: A report on glucose, oxidative stress and antioxidant biomarkers

**DOI:** 10.1101/2023.02.15.528625

**Authors:** Proteek Dasgupta

## Abstract

Fishes endemic to freshwater habitat are strongly influenced by water hardness with physiological consequences. The present study aimed to understand the effects of four-fold sequential increase from soft to hard waters, on selected tissues of Koi carp, a popular ornamental freshwater fish. Secondary stress markers - Glucose, Oxidative stress (Malondialdehyde – Lipid Peroxidation damage) and Antioxidants (Catalase, Glutathione-S-Transferase and Glutathione) were quantified in gills and white muscle after 14 days of exposure to hardness of 75 (Soft - TS), 150 (Moderate - TM), 225 (Hard - TH) to 300 (Very Hard - TV) mg CaCO_3_/L. Both the examined tissues were distinctly affected by soft and moderate waters. Glucose in gills (*p* < 0.05) was proportional to the rise in hardness concentration. Soft, moderate and very hard waters (75, 150 and 300 mg CaCO_3_/L) affected gills and muscle due to elevated MDA concentrations (*p* < 0.05). CAT and GST provided antioxidative protection to the tissues. The study results showed tissue-specific differential responses and more importantly, concentrations below 225 mg CaCO_3_/L elicited strong oxidative impairment in both gill and muscle.

## Introduction

Hardness is perhaps the most significant physicochemical property of water because it directly or indirectly influences physiology. Aquatic environments are constituted by varying concentrations of hardness, majorly due to Calcium (Ca^2+^) and Magnesium (Mg^2+^) along with some trace cations (Zn^2+^, Mn^2+^, etc) (Baldisserotto 2011; Romano et al. 2020). Based on the quantity of major cations, water is classified into soft (*<* 75 mg CaCO_3_/L) and hard (> 75 mg CaCO_3_/L) (Portz et al. 2006), both of which exert biological challenges. Soft water poses challenges to the survival of fish causing efflux across ion channels and destabilising ionic balance while, excessive hardness can cause hypercalcemia (Wendelaar Bonga et al. 1983) leading to bone ossification (Blanksma et al. 2009). It is, therefore, quite clear that water hardness leads to physiological changes. Without doubt, such changes can be studied through tissues which are largely impacted (Gonzalez et al. 1998; Gundersen and Curtis 1995).

Gill and muscle have provided substantial information about adaptation to water hardness in freshwater species such as Pinfish (*Lagodon rhomboides*) and Tilapia (Carrier and Evans 1976; Flik and Verbost 1995; Wendelaar Bonga et al. 1983). While gills are the epicentre of Ca^2+^ homeostasis and osmoregulation (Evans et al. 2005; Wendelaar Bonga et al. 1983), muscle is susceptible to changes in constituent amino acids and ionic shifts due to water hardness (Buentello and Gatlin 2002). Tendency of muscle mitochondria to take up Ca^2+^ from external sources and compensate for ionic stress has been reported in *Tilapia mossambica* (Sulochana et al. 1977). Both the tissues localise various redox reactions and such biomarker examinations provide information about physiological adjustments due to hard waters (Lushchak 2011).

Secondary stress markers are instrumental in assessing the effects of hard and soft waters. Reports by Copatti et al. (2019a) and Neves et al. (2017), highlight the usage of glucose to evaluate the effects of water hardness. Malondialdehyde (MDA) is a marker of lipid peroxidation (LPO), the consequences of which are prevented by enzymatic antioxidants (Catalase, Glutathione-S-Transferase) or non-enzymatic (Glutathione) thereby, rectifying the prooxidant/antioxidant ratio (Betteridge 2000; Lushchak 2016). Catalase is an important antioxidant enzyme that protects cells and tissues from oxidative damage because it reduces harmful hydrogen peroxide (H_2_O_2_) to water (H_2_O) (Betteridge 2000). Activity of GST is specific to detoxification of xenobiotics. It conjugates GSH to various electrophiles, thereby preventing oxidative damage. Although, GSH can also independently scavenge free radicals to defend the tissues from stress (Srikanth 2013). Despite broad insights offered by all the above, it is noteworthy that investigations involving its usage have gained momentum recently (Copatti et al. 2019b; Michelotti et al. 2018) to understand the extent and efficiency of physiological adaptations due to external hardness. Relevant information about the evaluation of secondary markers can make way for further molecular insights.

Koi carp, an ornamental variety of common carp, is popular among aquarists for aesthetical features. It is often spotted in various ponds, reservoirs, streams and lakes, which makes it an ecologically relevant species to study hardness and physiology interactions. Further, this species is physiologically sturdy under captive conditions as well. Therefore, the present study attempts to evaluate the effects of water hardness (75, 150, 225 and 300 mg CaCO_3_/L) on biomarkers (glucose, oxidative stress and antioxidant profile) in gill and muscle of Koi carp.

## Methods

### Acclimation of fish

Juveniles (6.70 ± 0.15 g; 5.90 ± 0.12 cm) were purchased from the Ornamental Fish Research Centre, Bengaluru. Fish were randomly distributed in 50 L glass tanks filled with tap water. The tanks were maintained under natural photoperiod (≈ 12 Light/12 Dark) with continuous aeration (Venus Aqua AP-608A, China). Fish were fed twice a day (09:00 and 18:00) at 2% body weight with commercial feed pellets (Taiyo Grow, India).

### Experimental setup

The two-week study consisted of four levels of hardness: 75 (Soft - TS), 150 (Moderate - TM), 225 (Hard - TH), and 300 (Very Hard - TV) mg CaCO_3_/L, selected based on its occurrence in natural systems (Stumm and Morgan 1996). Each treatment (hardness) was maintained in triplicate (6 fish per treatment). Hardness concentrations were prepared by the addition of Calcium carbonate (CaCO_3_) to water and calibrated by complexometric EDTA titration (APHA 2005). Faecal matter accumulated in the tank was drained out by suction pipe with subsequent renewal of the specific hardness. The tanks were covered with a mesh net to prevent the fish from escaping. Water parameters were monitored for temperature (25.1 ± 1°C), pH (7.04 ± 0.1), dissolved oxygen (6.5 ± 0.08 mg/L) and alkalinity (213 ± 0.02 mg/L) (APHA 2005).

### Sampling

Fish (*n* = 6 for each hardness level; total = 24 fish) were euthanized individually in clove oil solution (50 μL/L) and dissected for white muscle and gill. Tissues were washed with an ice-cold phosphate buffer solution (0.1M; pH 7.4) and homogenised in Potter-Elvehjem grinder. The homogenate (10% w/v) was then centrifuged at 5000 × *g* following which supernatant (stored at -20°C) was retained for all assays except Glutathione (homogenate precipitated with TCA before centrifugation). The absorbance values were measured on a spectrophotometer (Systronics UV-VIS 118, India).

### Biochemical analyses

#### Glucose

Glucose was assayed according to Nelson (1944) and Somogyi (1952). The deproteinizing agent (Ba(OH)_2_ and ZnSO_4_) was added to the supernatant and centrifuged at 5000 × *g* for 10 minutes. Alkaline copper reagent (potassium-sodium tartrate; Na_2_CO_3_; NaHCO_3_ and Na_2_SO_4_ in distilled water) was added to the supernatant. The mixture was heated, followed by the addition of an arseno-molybdate reagent. Optical density of the solution was recorded at 540 nm. A standard curve was plotted to evaluate the glucose concentration.

### Malondialdehyde (MDA)

Secondary product of Lipid peroxidation (LPO) - Malondialdehyde was estimated by using the Niehaus and Samuelsson (1968) protocol. The supernatant was mixed with this TCA-TBA-HCl reagent (15% Trichloroacetic acid, 0.38% Thiobarbituric acid and 0.25N Hydrochloric acid) in the ratio of 1:2. This reaction mixture was heated in a boiling water bath for 15 minutes, cooled and centrifuged at 1100 × *g* for 5 minutes. The optical density of the solution was recorded at 535 nm. MDA concentration was calculated using extinction coefficient of 1.56 × 10^5^ M^−1^ cm.

### Catalase (CAT)

Catalase activity was measured by using the Aebi (1984) protocol. The reaction was started by adding the supernatant to an equimolar solution of H_2_O_2_ and phosphate buffer (50 mM; pH 7.1). A decrease in absorbance was continuously recorded at 240 nm (UV) for 3 min.

### Glutathione-S-Transferase (GST)

GST activity was estimated by using the Habig et al. (1974) protocol. The reaction mixture contained the supernatant, phosphate buffer (0.1M; pH 6.5) and 2,4-Dinitrochlorobenzene (30mM). The volume was adjusted with distilled water after which the reaction was initiated by adding Glutathione (0.1 M). The optical density of the solution was recorded at 340 nm using a molar extinction coefficient of 9.6 × 10^3^M^−1^ cm^−1^.

### Glutathione (GSH)

GSH was estimated by using the Moron et al. (1979) protocol. Homogenate was precipitated with TCA (5%) and centrifuged at 3000 × *g* for 10 minutes. The supernatant collected after centrifugation was then added to the phosphate buffer (pH 6.5) and Ellman’s reagent. The optical density of the solution was recorded at 420 nm.

### Total protein

Total protein content of the tissues was estimated according to Lowry et al. (1951) protocol. Bovine serum albumin was used as standard. The optical density of the supernatant-reagents mixture was recorded at 660 nm.

### Statistical analysis

Data was summated as Mean ± SE. Normality and homoscedasticity were evaluated using the Shapiro–Wilk and Levene tests, respectively. Inter-group comparisons were performed using One-way ANOVA followed by a post-hoc test (Tukey). Significant differences were fixed at 95 % (*p* < 0.05). GraphPad Prism (Version 5.0, USA) and JASP (Version 0.16.2, Netherlands) were used for statistical computation and visual presentations.

## Results

### Biomarkers in Gill

The glucose concentration increased progressively from TS to TV. Significant differences (*F =* 10.91; *p* < 0.05) were found between TV and the remaining treatments (Figure 1A). Soft water TS elevated the concentration of MDA, followed by a spike in TV. Only TH differed significantly (*F =* 21.27; *p* < 0.001) from the remaining treatments (Figure 2A). Highest antioxidant Catalase activity was observed for TH, which differed significantly (*F =* 50.26; *p* < 0.001) from TS, TM and TV. Also, TV differed significantly from TM and TS (*F =* 50.26; *p* < 0.001) (Figure 3A). GST activity for TH and TV was comparatively higher than for TS and TM. While no significant differences (*F =* 26.45; *p* > 0.05) were found between the low (TS and TM) and high hardness groups (TH and TV), differences were observed between treatment pairs (Figure 4A). The highest concentration of GSH was recorded for TV. Except TS that was not significant (*F =* 42.78; *p* > 0.05) with TM and TH, remaining treatments recorded intergroup differences (Figure 5A).

**Figure 1.**
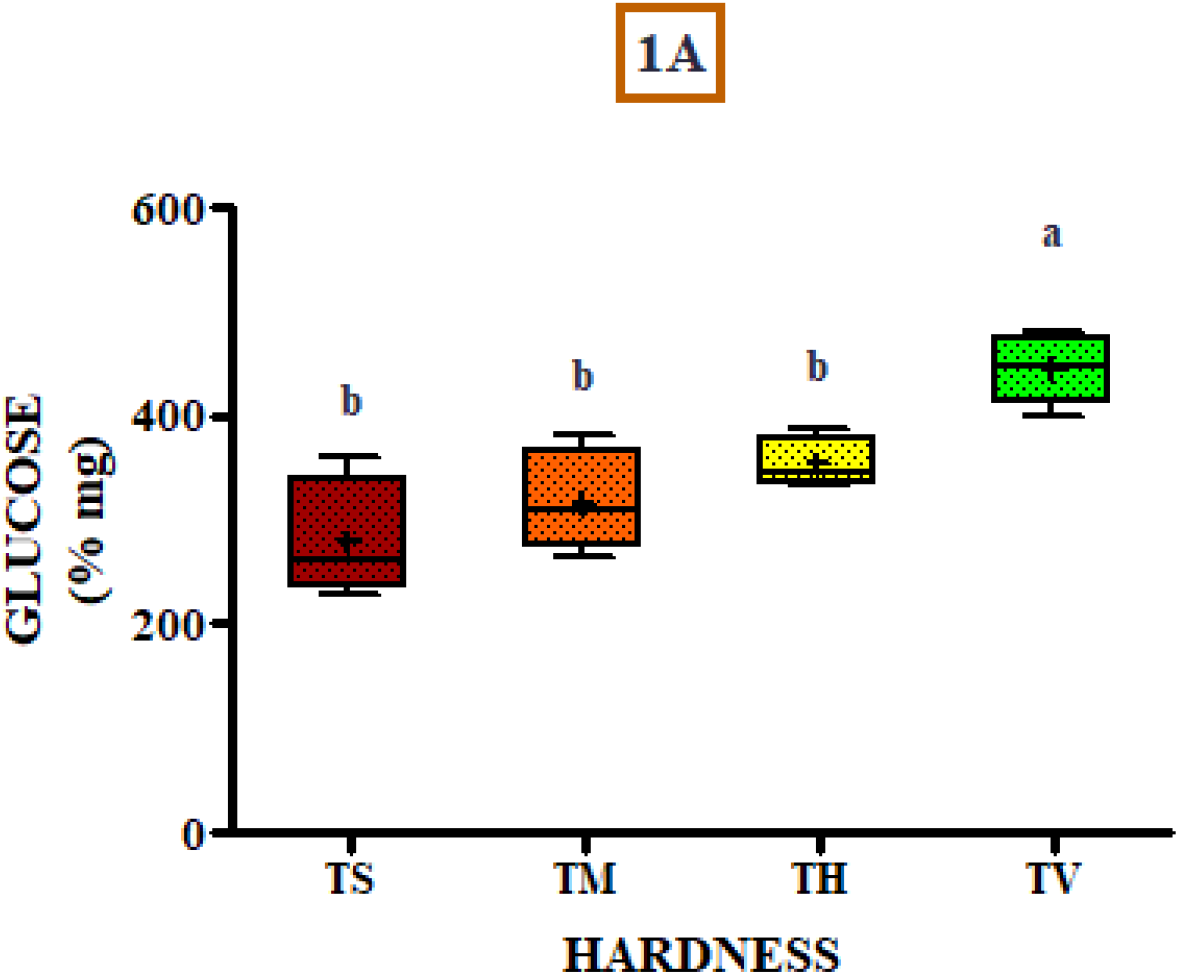
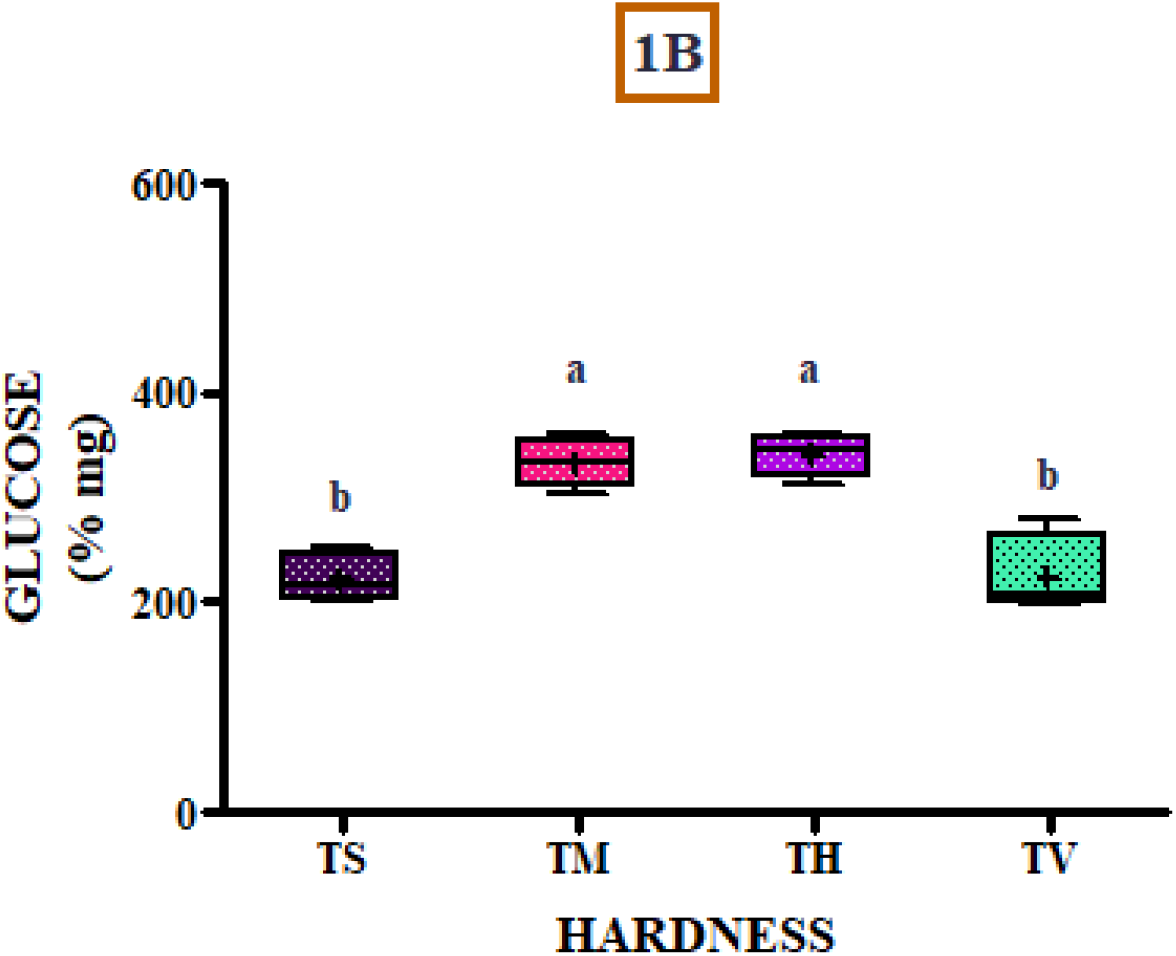
Effect of hardness concentration of 75 (Soft - TS), 150 (Moderate - TM), 225 (Hard - TH) and 300 (Very Hard - TV) mg CaCO_3_/L on glucose concentration in **(A)** gill and **(B)** white muscle of Koi carps (*n* = 24; 6 fish × 4 treatments). Superscripts among exposure groups indicate statistical significance (*p* < 0.05). (+) inside the box indicates the mean value of the exposure group.

**Figure 2.**
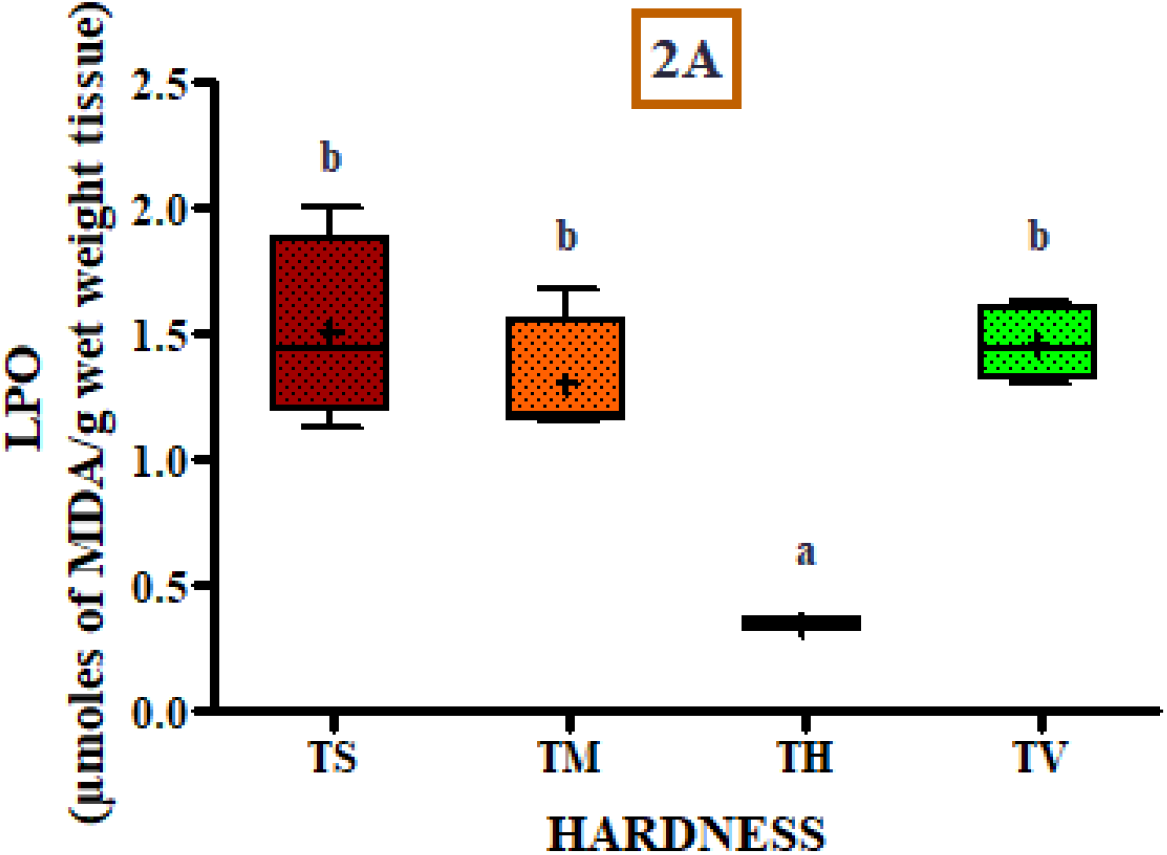
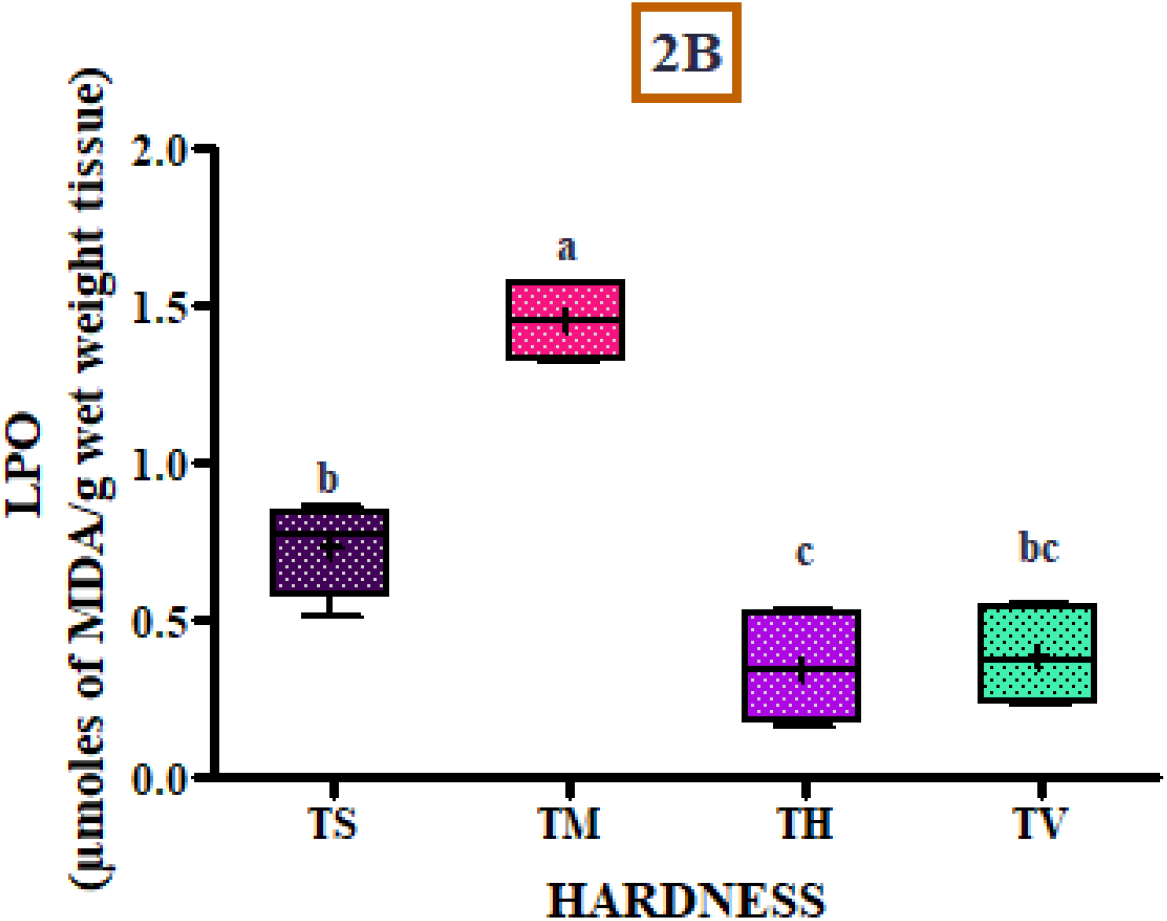
Effect of hardness concentration of 75 (Soft - TS), 150 (Moderate - TM), 225 (Hard - TH) and 300 (Very Hard - TV) mg CaCO_3_/L on MDA concentration in **(A)** gill and **(B)** white muscle of Koi carps (*n* = 24; 6 fish × 4 treatments). Superscripts among exposure groups indicate statistical significance (*p* < 0.05). (+) inside the box indicates the mean value of the exposure group.

**Figure 3.**
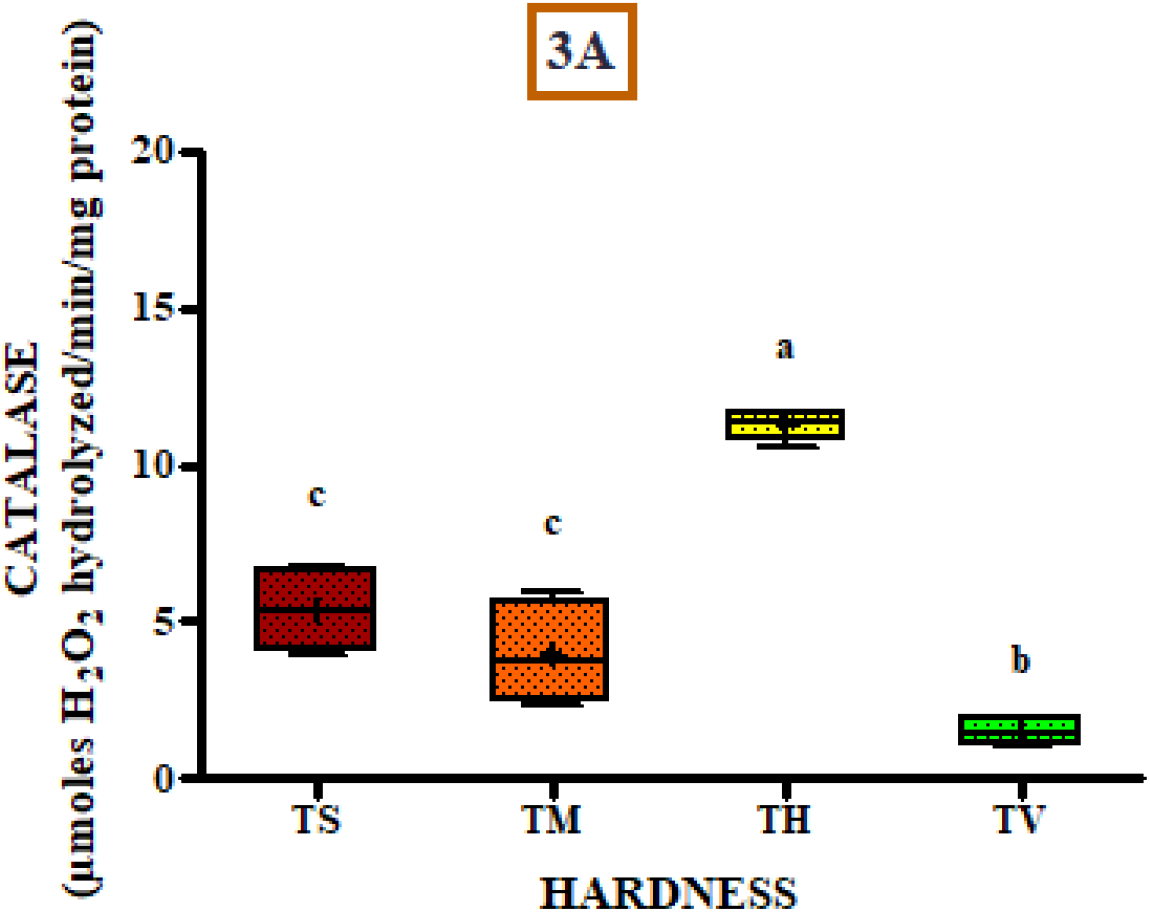
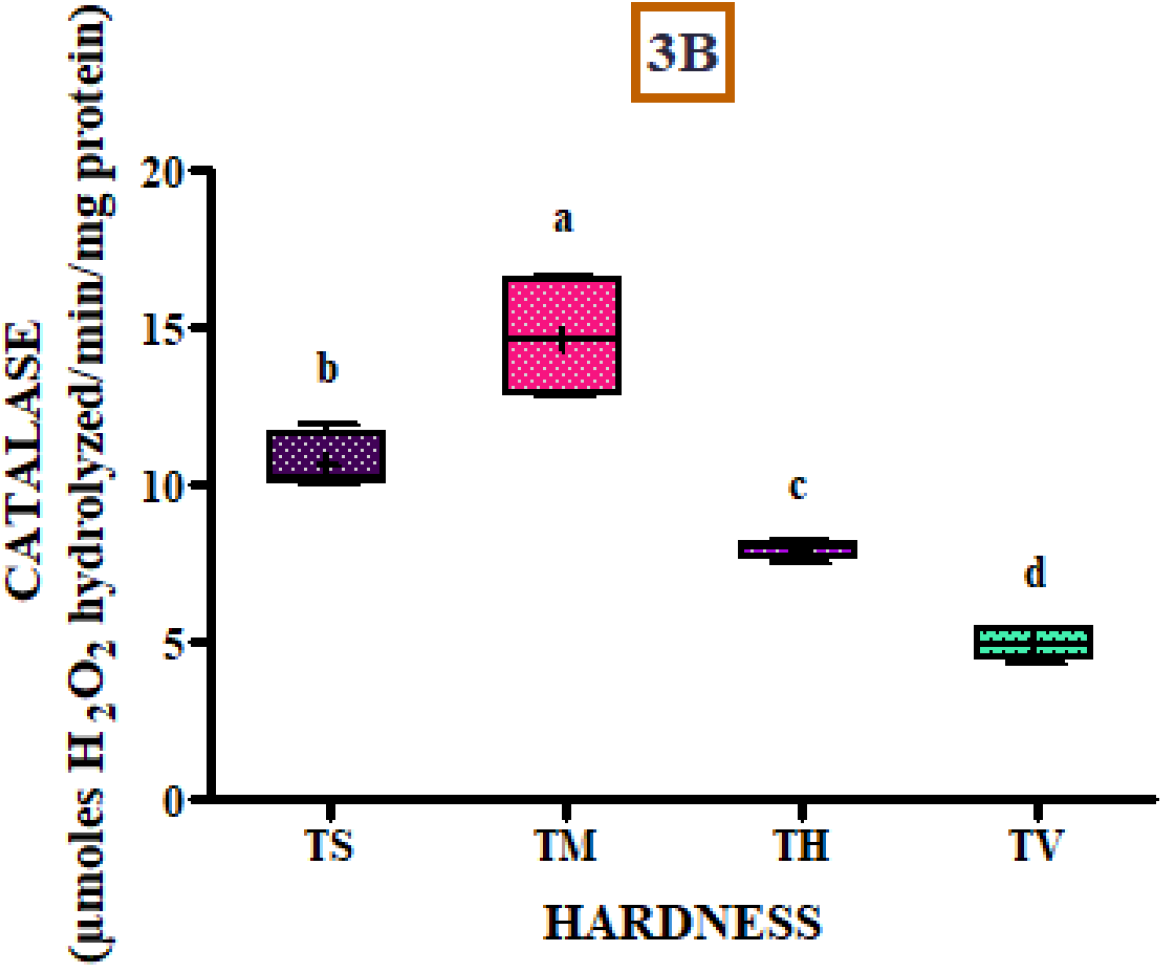
Effect of hardness concentration of 75 (Soft - TS), 150 (Moderate - TM), 225 (Hard - TH) and 300 (Very Hard - TV) mg CaCO_3_/L on Catalase activity in **(A)** gill and **(B)** white muscle of Koi carps (*n* = 24; 6 fish × 4 treatments). Superscripts among exposure groups indicate statistical significance (*p* < 0.05). (+) inside the box indicates the mean value of the exposure group.

**Figure 4.**
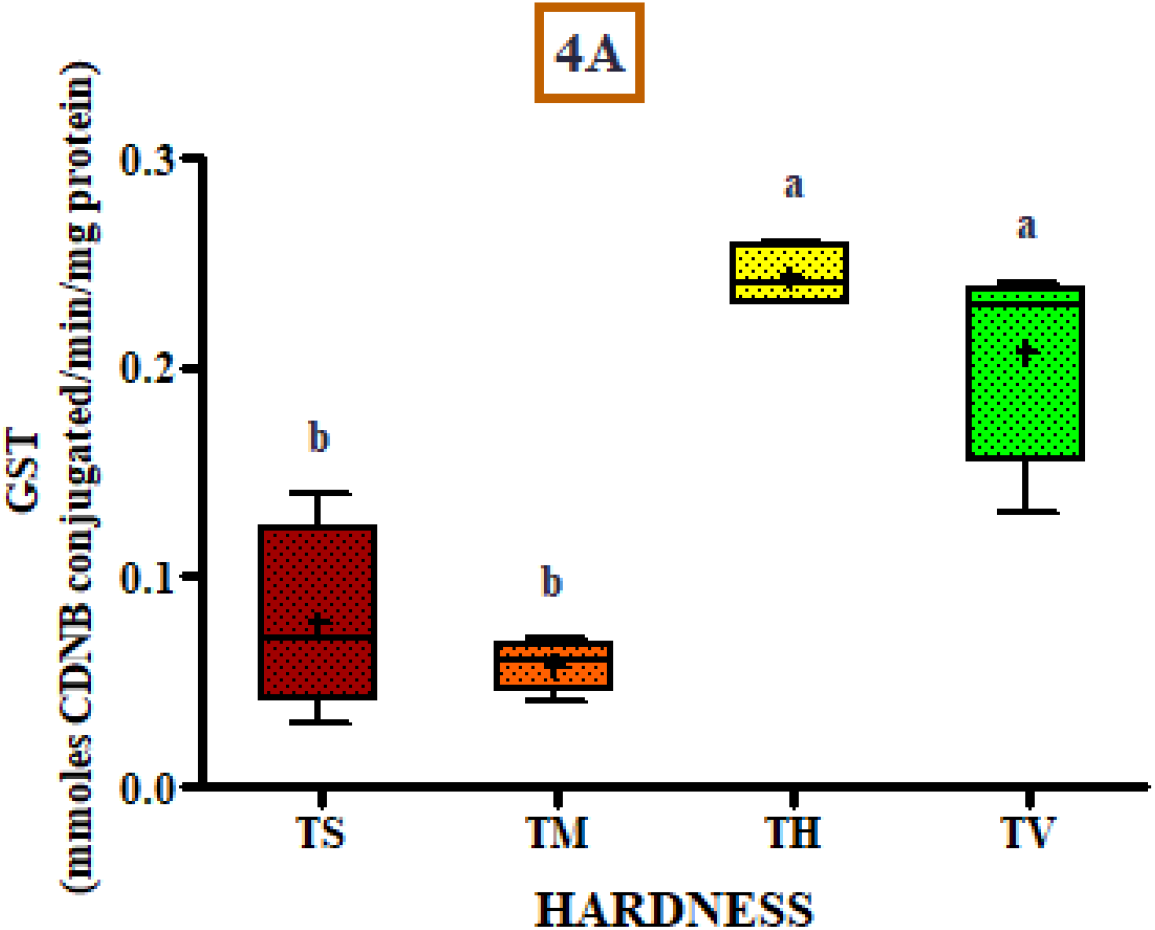
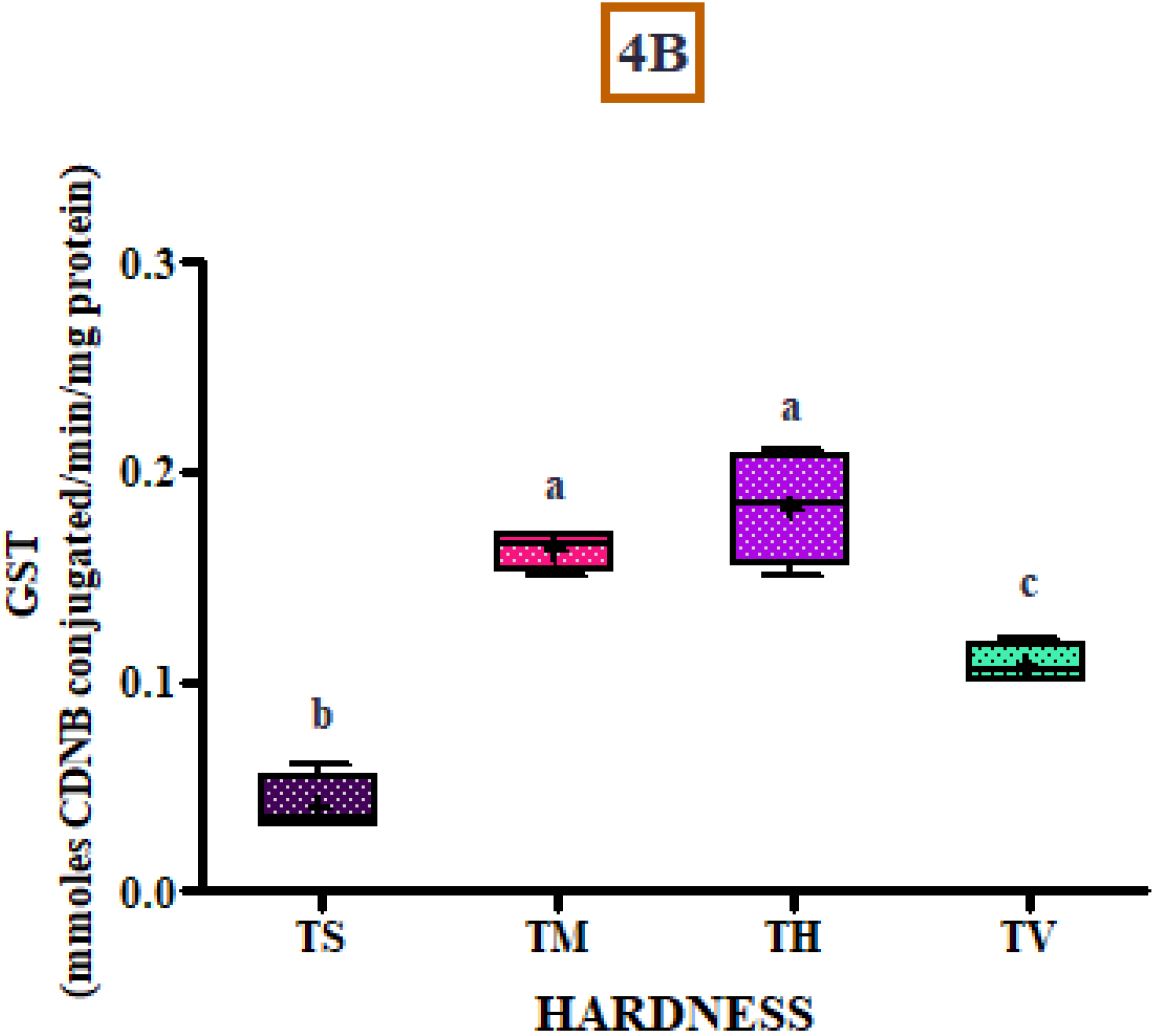
Effect of hardness concentration of 75 (Soft - TS), 150 (Moderate - TM), 225 (Hard - TH) and 300 (Very Hard - TV) mg CaCO_3_/L on Glutathione-S-transferase (GST) activity in **(A)** gill and **(B)** white muscle of Koi carps (*n* = 24; 6 fish × 4 treatments). Superscripts among exposure groups indicate statistical significance (*p* < 0.05). (+) inside the box indicates the mean value of the exposure group.

**Figure 5.**
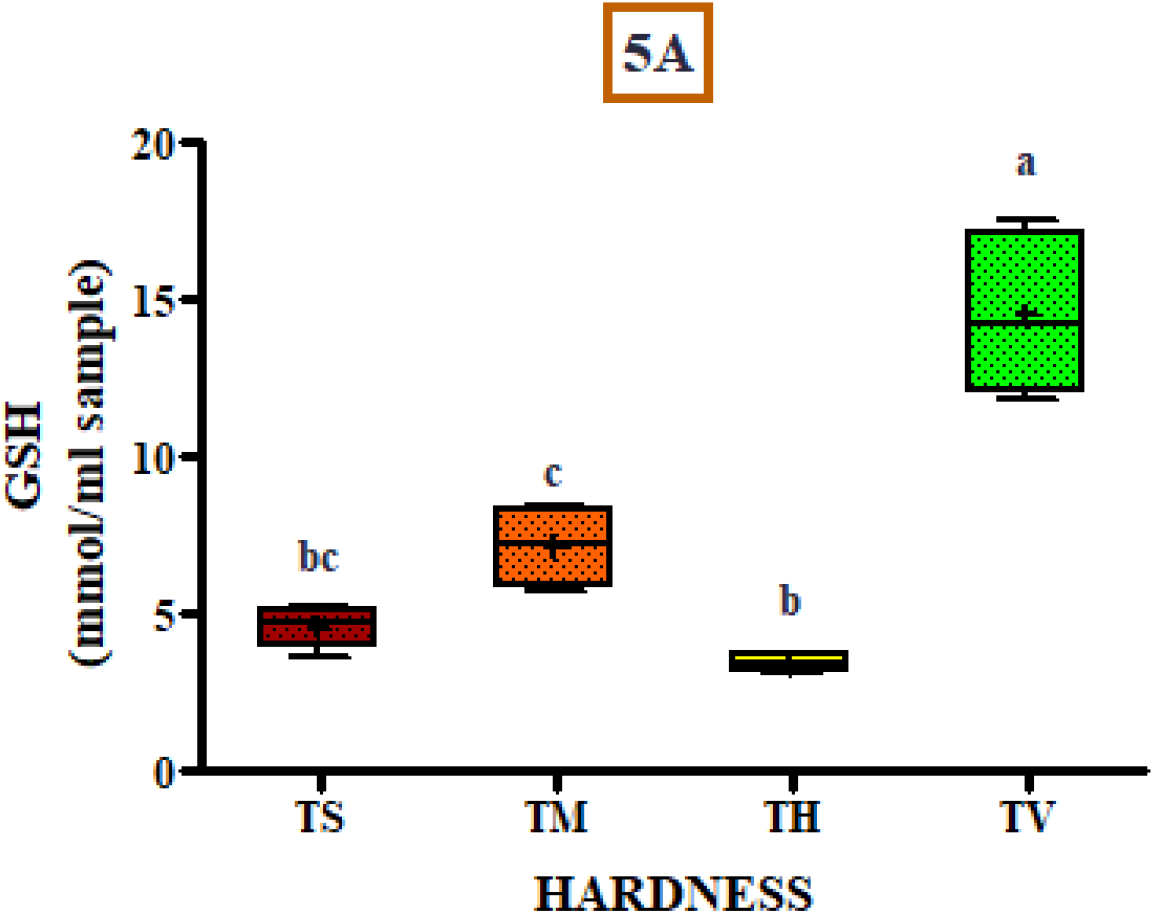
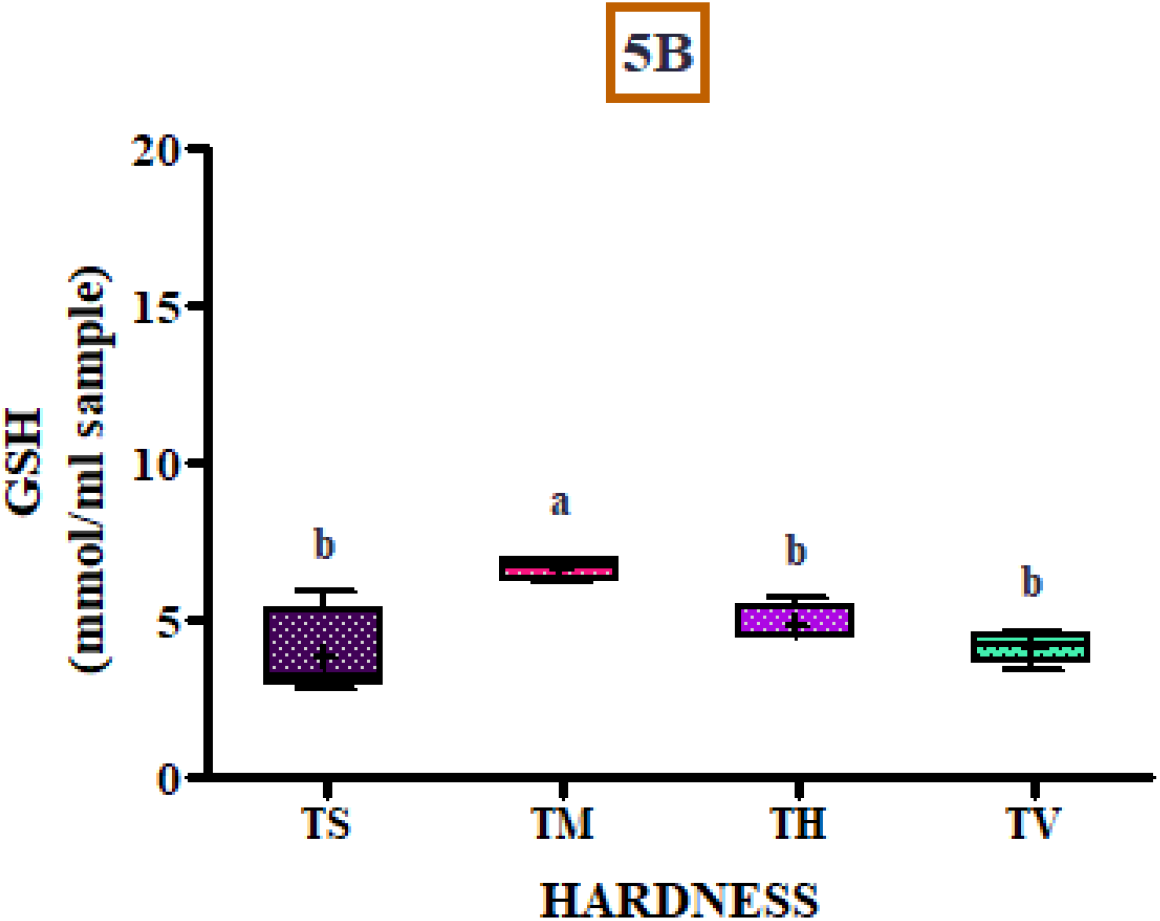
Effect of hardness concentration of 75 (Soft - TS), 150 (Moderate - TM), 225 (Hard - TH) and 300 (Very Hard - TV) mg CaCO_3_/L on Glutathione (GSH) in **(A)** gill and **(B)** white muscle of Koi carps (*n* = 24; 6 fish × 4 treatments). Superscripts among exposure groups indicate statistical significance (*p* < 0.05). (+) inside the box indicates the mean value of the exposure group.

### Biomarkers in White Muscle

The glucose concentration progressively increased from TS to TH but reduced at TV. Pairwise, similar concentrations and no significant differences (*F =* 22.98; *p* > 0.05) were observed for TS and TV and for TM and TH (Figure 1B). The highest level of MDA was observed for TM which differed significantly (*F =* 37.89; *p* < 0.001) from TS, TH and TV (Figure 2B). The highest CAT activity was observed in TM followed by a decrease for TH and TV. Catalase was the only biomarker that varied significantly (*F =* 49.38; *p* < 0.05) among all treatments for white muscles (Figure 3B). No significant differences (*F =* 57.13; *p* > 0.05) were recorded for GST between TM and TH. The highest GST activity was recorded for TH, which differed significantly with TS and TV (*F =* 57.13; *p* < 0.001) (Figure 4B). Glutathione (GSH) initially increased from TS to TM, thereafter, reducing gradually for the remaining treatments. Only significant difference (*F =* 8.87; *p* < 0.05) was observed between TM and the remaining groups (Figure 5B).

## Discussion

### Glucose concentration

In the present study, glucose increased sequentially in gills indicating that it was more conserved at higher levels of hardness. Progressive hardness led to increase in glucose, probably adding to the energy reserves. Since glucose serves as a primary energy for metabolism, glucose estimation can provide insights into the energy consumption for adaptations to hardness. Freshwater fish take up Ca^2+^ through the gills, and this transcellular movement is dependent on the surrounding Ca^2+^ concentrations, which affect the branchial permeability of the gills (Flik and Verbost 1995). Generally, an environment with high hardness reduces gill permeability (through tightening of cellular junctions) and subsequent loss of ions to water, ultimately conserving energy (Golombieski et al. 2013). This is clear through the results of the present study. Contrarily, muscle showed reductions in glucose at TV (300 mg CaCO_3_/L), which was also observed in juvenile Common Snook (*Centropomus undecimalis*) exposed to elevated hardness (Michelotti et al. 2018). Probably, increased energy demands for energy lowered muscle glucose levels, and it might have upregulated glycolysis.

### Oxidative stress

Presence of enormous amounts of Polyunsaturated Fatty Acids (PUFAs), predisposes fishes to lipid peroxidation, ultimately damaging cell membrane (Lushchak 2011) which is proportional to MDA produced. In the present study, excluding 225 mg CaCO_3_/L, MDA concentration for remaining exposures was elevated in gills. Possibly, soft (75), moderate (150) and very hard (300) waters, lead to damage of membrane associated with gills and white muscle. Subsequently, this could have led to excess production of MDA through free radical generation. In environments with low ionic concentration, certain membranes (such as apical membrane of gill) mechanise uptake of divalent cations from hard waters through Ca^2+^ channels embedded in it to meet the demand for necessary biological processes (Limbaugh et al. 2021). Elevated MDA is indicative of membrane damage and cell injury.

### Antioxidant response

Low catalase activity in gills exposed to soft (75) and moderate (30) waters indicated the failure to prevent oxidative damage. Low ionic composition of freshwater environment might burden the gills to reduce loss of ions to water for maintaining osmotic integrity. Therefore, efflux of ions to the external environment with concentrations *<* 150 mg CaCO_3_/L can ultimately lead to oxidative stress. Contrarily, muscle showed sequential decrease in antioxidant activity exposures above 150 mg CaCO_3_/L, largely remaining unaffected at higher concentrations.

Increase in GST activity was noticed in gills exposed to hard (225) and very hard waters (300). On the contrary, muscle showed elevated GST activity for all the exposures except soft waters (75). Though, higher GST activity clearly indicated greater antioxidant capacity in both the tissues, by far, antioxidative response was greater in muscle than gills. However, hardness *<* 75 mg CaCO_3_/L is detrimental for both the tissues and might impact osmoregulatory functioning (Evans 1987). Also, compared to Catalase, activity of GST was much higher in both the tissues for exposures above 225 mg CaCO_3_/L, indicating better antioxidant activity at higher levels of water hardness.

Antioxidant GSH was relatively low for muscle in contrast to gills, indicating better antioxidant capacity of muscle. In gills, a spike was observed for TH, which is conclusive that anything above 300 mg CaCO_3_/L is harmful. GSH can scavenge free radicals independently or in conjunction with GST to provide antioxidant defence (Srikanth et al. 2013). Clearly, the study results showed variance in GSH for both the tissues conforming to its specificity or tissue-specific antioxidant response. This has previously been reported in other popular freshwater species such as Nile Tilapia, Sharp Tooth Catfish (*Clarias lazera*) and Common carp (*Cyprinus carpio*) (Hamed et al. 2004).

## Conclusion

The study results reveal that soft and moderate (75 and 150 mg CaCO_3_/L) hardness can alter physiology of this freshwater species prompting antioxidative activity. Even after a prolonged period of 14 days, fish showed considerable adaptability to all exposures. Further, glucose concentration in gills was reserved for exposures to hardness > 225 mg CaCO_3_/L, to cope up in environments with low hardness. The results would benefit stakeholders and aquarist who culture this valued commercial species.

## List of abbreviations

MDA: Malondialdehyde
CAT: Catalase
GST: Glutathione-S-Transferase
GSH: Glutathione
GLU: Glucose
TS: Soft water
TM: Moderately hard water
TH: Hard water
TV: Very hard water

## Ethics approval

Animal use within this study was approved by the welfare committee of Bangalore University, India, in accordance with the national legislation.

## Availability of data and material

All relevant data is within this article. Please contact corresponding author for further queries

## Conflict of interest

The author declares that there are no known competing financial interests or personal relationships that could have appeared to influence the work reported in this paper.

## Funding

This research did not receive any grant from funding agencies in the public, commercial or not-for-profit sectors.

## Acknowledgements

Author thanks Bangalore University for providing necessary infrastructure.

## Notes

### Competing Interest Statement

The authors have declared no competing interest.

### Summary of Updates

Revision in Discussion section

